# RBPSignal: A deep learning approach for predicting RNA-Protein binding signals

**DOI:** 10.1101/2025.05.19.654811

**Authors:** Xiaohan Sally Ding, Zidong Alex Shu, Shengfan Wang, Luhao Shi, Yan Zhou, Biao Zhang, Hong-Bin Shen, Xiaojian Liu, Xiaoyong Pan

## Abstract

RNA-binding proteins play critical roles in post-transcriptional regulation by interacting with RNA to regulate various cellular processes. Inferring the binding signals and sequence patterns of these interactions is essential for elucidating the mechanisms underlying gene expression regulation. In this study, we present RBPSignal, a deep learning-based computational tool designed to predict RBP binding signals on RNAs and identify potential sequence motifs associated with these interactions. RBPSignal leverages the deep learning framework and trains on comprehensive eCLIP datasets, demonstrating an enhanced predictive accuracy. Furthermore, the integration of model interpretability through Integrated Gradients enables the detailed analysis of binding motif syntax. We validate the efficacy of RBPSignal on chromosome data and compare the discovered motifs with existing motif database, showcasing its ability not only to predict binding signals but also to uncover sequence patterns correlated with binding signals. Our findings provide insights into the sequence specificity of RBPs and explore the protein-protein interaction networks. RBPSignal serves as a valuable tool for exploring RBP-RNA interaction landscapes, facilitating further investigations into the regulatory networks underlying gene expression. The web server is freely available at http://www.csbio.sjtu.edu.cn/bioinf/RBPSignal/.

## Introduction

RNA-binding proteins (RBPs) are multifaceted regulators that control the fate of RNA post-transcriptionally, affecting their splicing, stability, translation and degradation(Khoroshkin et al., 2024). Abnormal binding of RBPs to RNAs has been implicated in the pathogenesis of various diseases, including cancer, neurodegenerative disorders and developmental abnormalities(Gebauer et al., 2021; Kelaini et al., 2021; Nutter et al., 2016; Wang et al., 2022). Understanding the intricate interactions between RBPs and RNAs is becoming increasingly crucial for unraveling the complex regulatory networks within cells. The binding specificity of RBPs to RNAs is determined by a combination of factors, including the sequence, structure, and chemical modifications of the RNA molecules(Verta & Jacobs, 2024). Therefore, elucidating the binding signals and sites of RBPs on RNAs is vital for comprehending the binding mechanisms and enabling subsequent analyses.

Recent developments in CLIP-derived technologies have facilitated the precise identification of RBP binding sites at base resolution. Key techniques in this field, including Photoactivatable-Ribonucleoside-Enhanced CLIP (PAR-CLIP)(Hafner et al., 2010), Individual-Nucleotide-Resolution CLIP (iCLIP)(König et al., 2010) and Enhanced CLIP (eCLIP)(Van Nostrand et al., 2016), provide enhanced sensitivity and specificity for detecting RBP-RNA interactions, thereby deepening our understanding of the regulatory roles in various biological processes. Encyclopedia of DNA Elements (ENCODE) project has utilized eCLIP combined with high-throughput sequencing (eCLIP-seq) to gather comprehensive binding sites for RNA-binding proteins in the HepG2 and K562 cell lines, and in adrenal gland tissue(Moore et al., 2020). By employing eCLIP-seq, RNA crosslinking sites can be accurately located, which aids in identifying RBP-binding RNAs and elucidating binding motifs.

Currently, various computational tools have been developed to predict the binding sites of RBPs on RNAs, essential for understanding their roles in gene expression regulation and disease mechanisms. For instance, DeepBind applies convolutional neural networks to accurately predict the sequence specificities of DNA- and RNA-binding proteins across diverse datasets(Alipanahi et al., 2015). RNAcompete measures RBP binding specificity to RNAs and generates training data for DeepBind models(He et al., 2016). DeepSite(Jiménez et al., 2017) leverages a 3D deep convolutional neural network to identify the binding site using protein structures. Notable tools include RBPsuite, which uses deep learning to predict RNA-protein binding sites for both linear and circular RNA(Pan et al., 2020). PrismNet(Sun et al., 2021) was developed to accurately predict dynamic RBP binding across various cellular conditions by integrating in vivo RNA structure data. Additionally, Zhu et al. proposed HDRNet(Zhu et al., 2023), an end-to-end deep learning model for identifying RNA-binding interactions from eCLIP-seq data across cellular conditions.

However, despite the advancements, many models still focus on classification tasks and rely on datasets that carry biases from CLIP-seq experiments(Horlacher et al., 2023). For example, the widespread use of 1:1 positive-to-negative sample ratios and ambiguous definitions of negative events can misrepresent actual biological distributions, resulting in false-positive predictions for full-length RNA(Sun et al., 2021; Wu et al., 2021). Addressing these challenges requires the development of more refined dataset and advanced regression models capable of capturing RNA sequence context and sequential characteristics with greater precision.

Here, we developed RBPSignal, which is designed to predict the binding signals between RBPs and RNAs using a regression model, while also identifying potential binding sequence patterns associated with the interactions. Beyond signal prediction, RBPSignal enables the identification of putative binding motifs-sequence patterns that contribute to RBP-RNA interactions. To achieve this, RBPSignal leverages the eCLIP dataset, incorporating a sliding-window approach during data preprocessing to systematically segment full-length transcripts, ensuring comprehensive sequence coverage and contextual integrity. This preprocessing strategy not only enriches training data diversity but also facilitates finer resolution in detecting binding signal variations along transcripts. RBPSignal model architecture integrates a two-layer of convolutional neural network (CNN), a bidirectional long short-term memory (BiLSTM) network, and a three-layer linear module to capture RBP binding sequence patterns and model interpretability through Integrated Gradients (IG). Unlike previous classification-based models such as RBPBind(Gaither et al., 2022) and GraphProt(Maticzka et al., 2014), RBPSignal focuses on the regression-based task, leveraging the eCLIP datasets’ unique inclusion of control groups to comprehensively extract and utilize informative signals. With exceptional performance in both prediction accuracy and interpretability, RBPSignal bridges the gap between binary classification with continuous regression tasks and facilitates the study of RBP-motif-RNA networks through an online web server, providing a user-friendly platform for predicting RBP binding signals and promoting further exploration of their biological significance.

## Method and materials

### Overview of RBPSignal

RBPSignal is designed to predict the binding signals of RNA-binding proteins on RNA sequences. A novel data processing method is employed to prepare datasets for model training. Unlike traditional ENCODE methodologies, which truncate sequences to 101 nt length centered at the nucleotide position that holds the peak signal (50 nt upstream and 50 nt downstream), RBPSignal applies a sliding-window approach across the entire length of transcripts, with a sliding step of 30 base pairs (Fig. 1a). This approach enables the model to capture continuous signals along RNA sequences and detect changes in binding patterns. The specific processing principles are illustrated in Fig. 1b. The workflow begins with BAM file processing through htseq-clip to extract RNA-binding protein crosslinking signal, followed by a sliding window to segment the crosslinking signal into fixed-length windows. The crosslinking signal of sliding windows is then analyzed using DEWseq to identify differentially enriched binding windows, filtering out non-significant interactions considered as ‘noise’ while retaining both high-confidence binding and non-binding regions. Finally, sequences of binding regions are extracted via subseq for subsequent analysis including as model input, motif discovery, Gene Ontology (GO) enrichment studies, ensuring a systematic and reliable identification of functional RBP binding sites.

**Figure 1.**
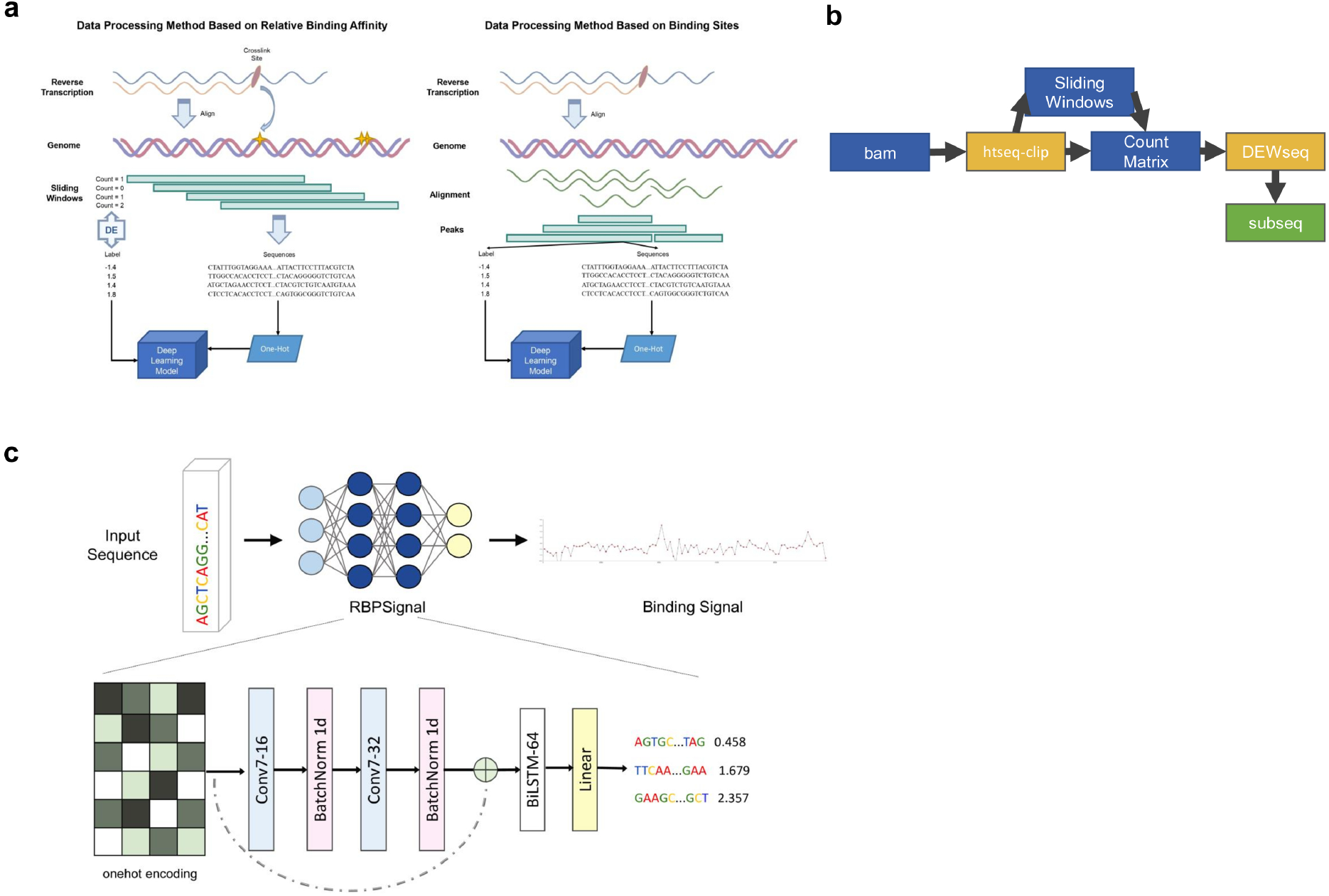
Workflow and data processing of RBPSignal. a. Comparative analysis of data processing between RBPSignal and traditional ENCODE methodologies. b. Illustration of the RBPSignal data processing pipeline c. Overview of RBPSignal.

RBPSignal consists of a convolutional neural network (CNN), a bidirectional Long Short-Term Memory network (BiLSTM), and a multilayer perceptron (MLP) block. The two-layer CNN is designed to capture regional relationships along the RNA sequence, while incorporating residual connections to mitigate information loss (Fig. 1c). The CNN output is then processed by a BiLSTM, which captures sequential dependencies and contextual relationships in both directions. Finally, the MLP block reduces the dimensionality of the output, ensuring that each sequence fragment corresponds to a single output value, representing the RBP binding signal.

### Benchmark dataset collection

RBPSignal utilizes eCLIP data from the ENCODE dataset, which is sourced from the following link: https://www.encodeproject.org/search/?type=Experiment&status=released&internal_tags=ENCORE&assay_title=eCLIP&biosample_ontology.term_name=K562&biosample_ontology.term_name=HepG2. A key advantage of eCLIP data is the inclusion of size-matched input (SMI) control samples, which are processed in parallel with immunoprecipitation (IP) samples. These controls are essential for accounting for background signal and allow for more accurate discrimination of specific binding events. For our project, we selected 165 high-quality eCLIP datasets that provided sufficient sequencing depth from eCLIP dataset with a total of 252 consistently generated experimental studies. These datasets correspond to 124 unique RBPs across two human cell lines, HepG2 and K562; 1 tissue cell line: adrenal gland. Our selection criteria required that each RBP be represented by at least two independent biological replicates to capture consistent binding patterns and minimize the impact of experimental variability.

### Genome annotation profile

We mapped CLIP-derived crosslink sites to their corresponding target genes using the GENCODE comprehensive gene annotation for the human genome (GRCh38, release 41). This annotation file facilitated the alignment of our sequencing data with the human genome, enabling precise localization of RNA-protein interactions within known genomic features.

### Data processing for model training

In terms of data processing, we initially employed a Python software package named htseq-clip(Sahadevan et al., 2023) for the statistics of crosslinking sites. This package is specifically designed for preprocessing, extraction, and summarization of crosslinking site counts from iCLIP and eCLIP sequencing experiment data. We used hg38 as the reference genome and the alignment files as inputs which were sourced from the ENCODE eCLIP dataset. htseq-clip allowed us to generate a sliding window of length 101 nucleotides and segment each gene annotation feature with a step size of 30 nucleotides. For each segment, the tool provided a count of crosslinking sites, and these counts were summarized in a matrix format.

Then, we integrated the use of DEseq2 and DEWseq software packages for differential expression analysis between the experimental and control groups. The method of processing eCLIP data with DEWseq has been mentioned in previous studies(Schwarzl et al., 2024). We utilized DESeq2’s functions during differential expression analysis to filter out non-significant regions while retaining high-confidence positive and negative signal samples.

Subsequently, we retrieved the genomic sequences corresponding to the sliding windows using bedtools, a tool designed to extract nucleotide sequences based on genomic coordinates. The sequences extracted from the sliding windows were then matched with the signal values obtained from the differential expression analysis.

### Motif database preparation

We employed the combined collection of motifs from the MEME(Bailey et al., 2015) and DEWSeq(Schwarzl et al., 2024) study for motif annotation analysis. First, the relevant motif files from these databases were downloaded, which contain position weight matrices (PWMs) for various RNA-binding proteins. This merged motif database, formatted in MEME format, then used to evaluate motif discovery results from RBPSignal. Rather than relying on prior motifs for training, RBPSignal first identifies sequence patterns associated with high binding signal. After discovering putative motifs from the model’s learned sequence features, these motifs were compared against this database.

### RBPSignal parameter settings

The model comprises three primary components: a convolutional neural network block, a long short-term memory (LSTM) layer, and a linear output layer:

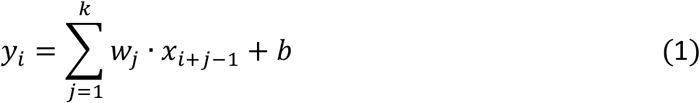

*x* ∈ *R*^*C*×*L*^ represents the input sequence with *C* channels (e.g., *C* = 4 for one-hot encoded nucleotide bases A, C, G, and T) and sequence length *L*. The convolutional filter is denoted as *w* ∈ *R*^*C*×*L*^, where *k* is the kernel size, and the filter spans all input channels. The output feature map at position *i* is given by *y*_*i*_, and *b* ∈ *R* is a scalar bias term. The index *j* refers to kernel position (ranging from 1 to *k*) and the term *i* + *j* − 1 defines the location in the input sequence currently being convolved with RELU activation:

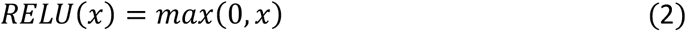

Specifically: for positive inputs (*x* > 0), ReLU returns the input itself *x*; negative inputs (*x* ≤ 0), ReLU outputs 0. This non-linear transformation introduces sparsity in the neural network which can be beneficial for both model efficiency and training performance.

The loss function is defined as:

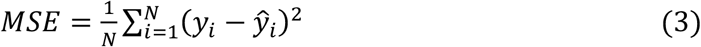

where N represents the number of samples in a batch, *y*_*i*_ is the true signal for the *i*-th sample, and *ŷ*_*i*_ is the corresponding prediction value generated by the model.

The CNN block begins with a 1D convolutional layer with 16 filters, each of size 7, and padding of 3 to preserve sequence length. This is followed by batch normalization and ReLU activation. A second convolutional layer with 32 filters (also kernel size 7 and padding 3) further processes the features, with an additional batch normalization layer to stabilize training. A residual connection integrates the input with the output, using a 1D convolutional downsampling layer (1×1 kernel). The processed features are passed to the LSTM layer, which employs a single-layer LSTM with an input size of 32 (matching the output channels of the CNN block) and a hidden state size of 64. Finally, the fixed-size representation is the linear layer module, consisting of three fully connected layers. The first dense layer reduces the input from 64 to 128 hidden units, followed by ReLU activation and a dropout layer (rate 0.4) to prevent overfitting. The second dense layer further reduces the representation to 64 hidden units with another ReLU activation. The final dense layer produces a single output representing the predicted binding affinity.

### Motif discovery through Integrated Gradients

To identify motifs associated with RBPs binding, we utilized Integrated Gradients (IG) to calculate the contribution score for each base within 101 bp length sequences. A sliding window of size 7 was applied to identify the highest-scoring motifs for each sequence. Subsequently, sequences were ranked based on their predicted binding scores, and the top 40,000 sequences across all datasets were selected for motif analysis. If fewer than 40,000 sequences were available, all sequences were included. The formula of IG is as follow:

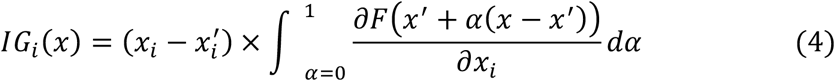

where x denotes the actual input sequence, *x’* represents a baseline input (a sequence of zeros in our calculation), F is the output of the predictive model, and *α* is a scalar that interpolates between the baseline and input. The integration aggregates gradients along the linear path from *x’* to *x* attributing a contribution score to each base *x*_*i*_. The continuous gradient-based evaluation provides a principled method for interpreting deep learning models in biological sequence analysis.

To annotate the identified motifs with known databases, we used the FIMO, a tool for scanning sequences for occurrences of known motifs, to identify matching patterns of combined motif collection. Motifs that were not found in the MEME file underwent further analysis, with the two most frequently occurring motifs selected to represent the corresponding RBP file.

### Comparison of RBPSignal with other methods

To evaluate the performance of RBPSignal, we compared it against RBPsuite and DeepBind using mitochondrial chromosome data. The evaluation involved calculating the average Spearman correlation coefficient (SCC) and the maximum area under the curve (AUC).

For AUC calculation, the continuous binding signals were first binarized by applying a series of uniformly spaced thresholds between the minimum and maximum signal values. Specifically, 20 thresholds were generated across this range. At each threshold, fragments with signals greater than or equal to the threshold were labeled as 1 (positive binding), and those below as 0 (negative binding). AUC was then computed between the predicted continuous outputs and these binarized labels, with the highest AUC across all thresholds reported as the final score. Each RBP file contains multiple transcripts, which were segmented into 101-nt fragments. We calculated the maximum AUC for each transcript and then averaged these values across all transcripts. The comparison across the three tools was conducted on 27 specific RBPs that were commonly available for prediction across all methods, including FUS_K562, FXR1_K562, GTF2F1_HepG2, GTF2F1_K562, HNRNPA1_HepG2, HNRNPA1_K562, HNRNPC_HepG2, HNRNPK_HepG2, HNRNPK_K562, HNRNPL_HepG2, IGF2BP2_K562, IGF2BP3_HepG2, KHDRBS1_K562, MATR3_HepG2, MATR3_K562, PABP4_K562, PCBP1_HepG2, PCBP2_HepG2, QKI_HepG2, QKI_K562, SRSF1_HepG2, SRSF1_K562, SRSF7_HepG2, SRSF7_K562, SRSF9_HepG2, TIA1_K562, and U2AF2_HepG2.

### Protein-protein interactions and Gene Ontology (GO) analysis

To investigate protein-protein interactions, we hypothesized that RBPs sharing similar binding motifs may belong to the same protein complex, potentially facilitating their interactions. After motif discovery, we mapped all identified motifs of each RBP to a motif database, identifying proteins that share the same motifs. We then selected RBPs with a q-value < 0.05 as candidate interacting proteins. By iterating this process, we constructed a putative protein-protein interaction network. To validate our network, we compared it with the STRING database (Version 12.0)(Szklarczyk et al., 2023).

For phylogenetic analysis, we used MEGA (Version 11)(Tamura et al., 2021) with ClustalW to perform multiple sequence alignments of all identified motifs. If an RBP’s motifs were mapped to the database, those motifs were used; otherwise, the top two most frequent motifs were selected. A Neighbor-Joining tree was then constructed. Gene annotation and enrichment analysis were conducted using Metascape(Zhou et al., 2019), selecting functional categories potentially associated with motif phylogeny for RBP annotation. Finally, the phylogenetic tree and functional annotations were visualized using iTOL (Version 6.0)(Letunic & Bork, 2024).

## Results

### RBPSignal outperforms baseline methods

To objectively evaluate the performance of RBPSignal and compare it with traditional methods, we constructed benchmark datasets derived both from our RBPSignal data generation pipeline and from ENCODE-generated processing. The data was split based on chromosomes to avoid any overlap between training and testing samples. Chromosomes 2, 9, and 16 were selected for validation, while chromosomes 1, 8, and 15 were used exclusively for testing. All other chromosomes, excluding the mitochondrial chromosome, were allocated to the training dataset.

Performance metrics include the Spearman correlation coefficient (SCC) and Area Under the ROC Curve (AUC). We compared the ranking consistency of binding predictions using Spearman correlation coefficient (SCC) across transcripts, and assessed classification performance using the average of the maximum AUC values per transcript. For each RBP file, we averaged these transcript-level scores to obtain a single representative value per RBP. As shown in Fig 2a and 2b, the model trained on the RBPSignal-generated test dataset outperformed the one trained on the ENCODE test dataset under both evaluation criteria.

**Figure 2.**
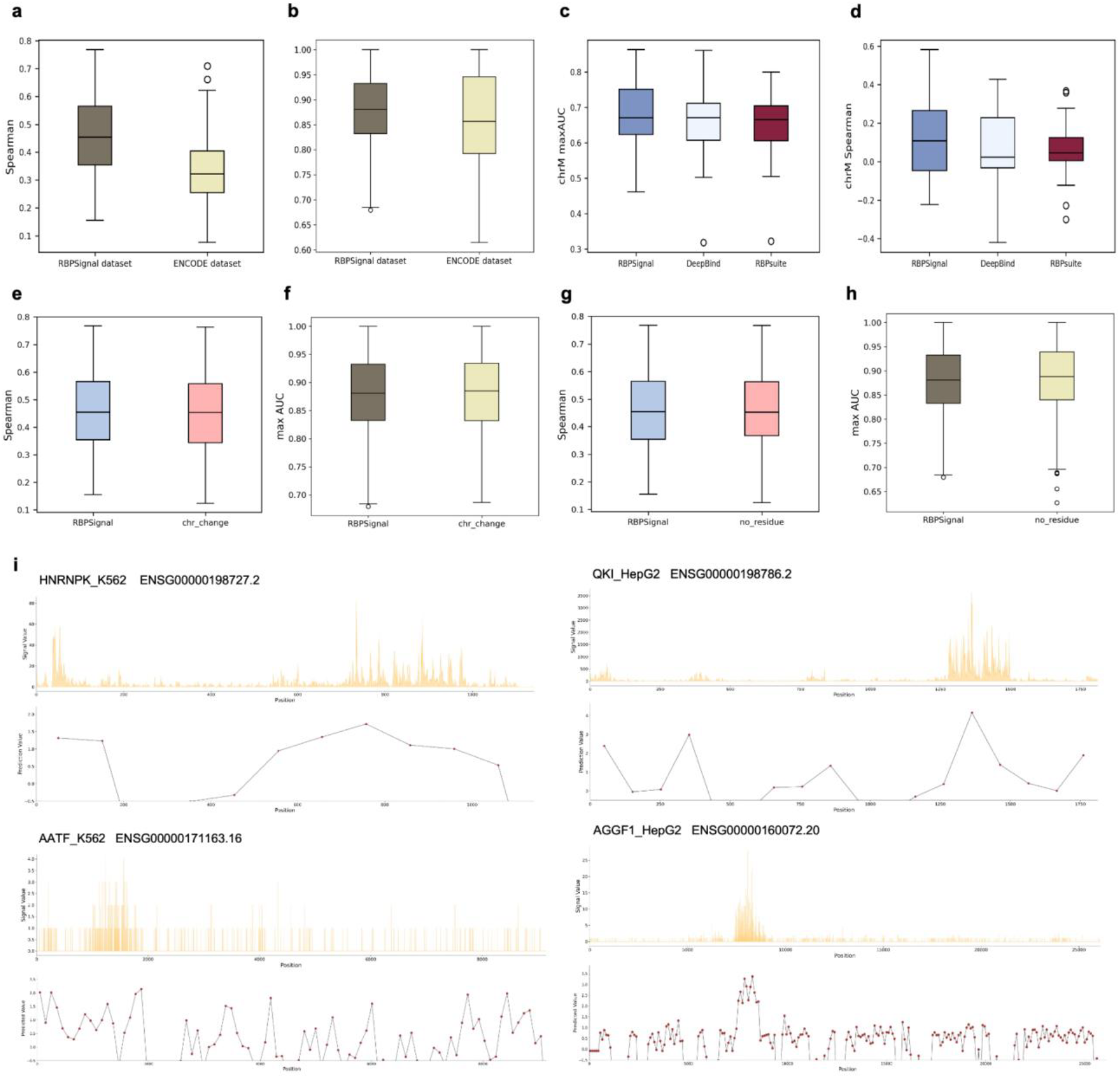
RBPSignal achieves superior performance in signal prediction on test datasets and mitochondrial chromosomes. a. Comparison of Spearman correlation coefficients for model training on RBPSignal datasets versus ENCODE datasets. b. Comparison of maximum AUC values for model training on RBPSignal datasets versus ENCODE datasets. c. Spearman correlation coefficient comparison among RBPSignal, DeepBind, and RBPsuite for mitochondrial chromosome predictions. d. Maximum AUC comparison among RBPSignal, DeepBind, and RBPsuite for mitochondrial chromosome predictions. e. Spearman correlation coefficients comparison with chromosome dataset shifting. f. Maximum AUC comparison with chromosome dataset shifting. g. Spearman correlation coefficients comparison with model variant without residue block. h. Maximum AUC comparison with model variant without residue block. i. Visualization of different RBP binding signals.

Additionally, existing methods such as DeepBind and RBPsuite, which predict binding scores for individual RNA sequence fragments, were compared to RBPSignal. SCC and maximum AUC values were calculated on mitochondrial chromosome data, revealing that RBPSignal achieved superior predictive performance compared to these methods (Fig. 2c and 2d).

To further validate the robustness and architectural design of RBPSignal, we conducted an ablation study from two complementary perspectives: (1) the impact of chromosome reassignment in data partitioning, and (2) the contribution of residual blocks in the model architecture. This dual strategy ensures that both the input data division and network design do not introduce artifacts or overfitting, and that RBPSignal generalizes effectively across genomic contexts and structural depths. For the first strategy, we reassigned the validation and test datasets to new chromosome sets. While the original validation chromosomes were 2, 9, and 16, and the test chromosomes were 1, 8, and 15, in this variant setting, chromosomes 3, 10, and 17 were used for validation, and chromosomes 4, 11, and 18 for testing. The training chromosomes remained unchanged (excluding mitochondrial and those used for validation and test). This ensured non-overlapping training and test sets, thereby avoiding data leakage. Performance was evaluated on both the original and reassigned test sets using SCC and AUC. The results showed no significant deviations in SCC or AUC before and after chromosome reassignment (Fig. 2e and 2f).

When evaluated on the same original test set, the model variant without residual connections exhibited greater variance in performance across different RBPs, including three extreme outliers, compared to only one in the original RBPSignal model (Fig. 2g and 2h). This suggests that residual block contributes to more stable learning by facilitating better information integration across layers, helping the model avoid information degradation.

Examples of RBPs HNRNPK (K562), QKI (HepG2), AATF (K562), and AGGF1 (HepG2), with corresponding transcripts-ENSG00000198727.2, ENSG00000198786.2, ENSG00000171163.16, and ENSG00000160072.20, are illustrated in (Fig. 2i). RBPSignal successfully predicted the binding signals on both long and short transcripts for both cases, showing the potential binding signal values of each 101-nt RNA sequence with positive values displayed.

### Discovering motifs and providing novel biological insights into RBPs

RNA-based RBP binding motifs play a critical role in mediating RBP-RNA interactions, and unraveling these binding motifs provides deeper insights into the underlying binding mechanisms. To explore these interations, we applied Integrated Gradients method to calculate the contribution score of each base within RNA sequences. As examples shown in Fig. 3a, sequences with motifs substantiated in existing databases (a merge of Ray2013 and catRAPID databases) for DD3X_HepG2, QKI_HepG2, and RBFOX2_K562 illustrate that the RBPSignal model effectively captures highly-contributive bases.

**Figure 3.**
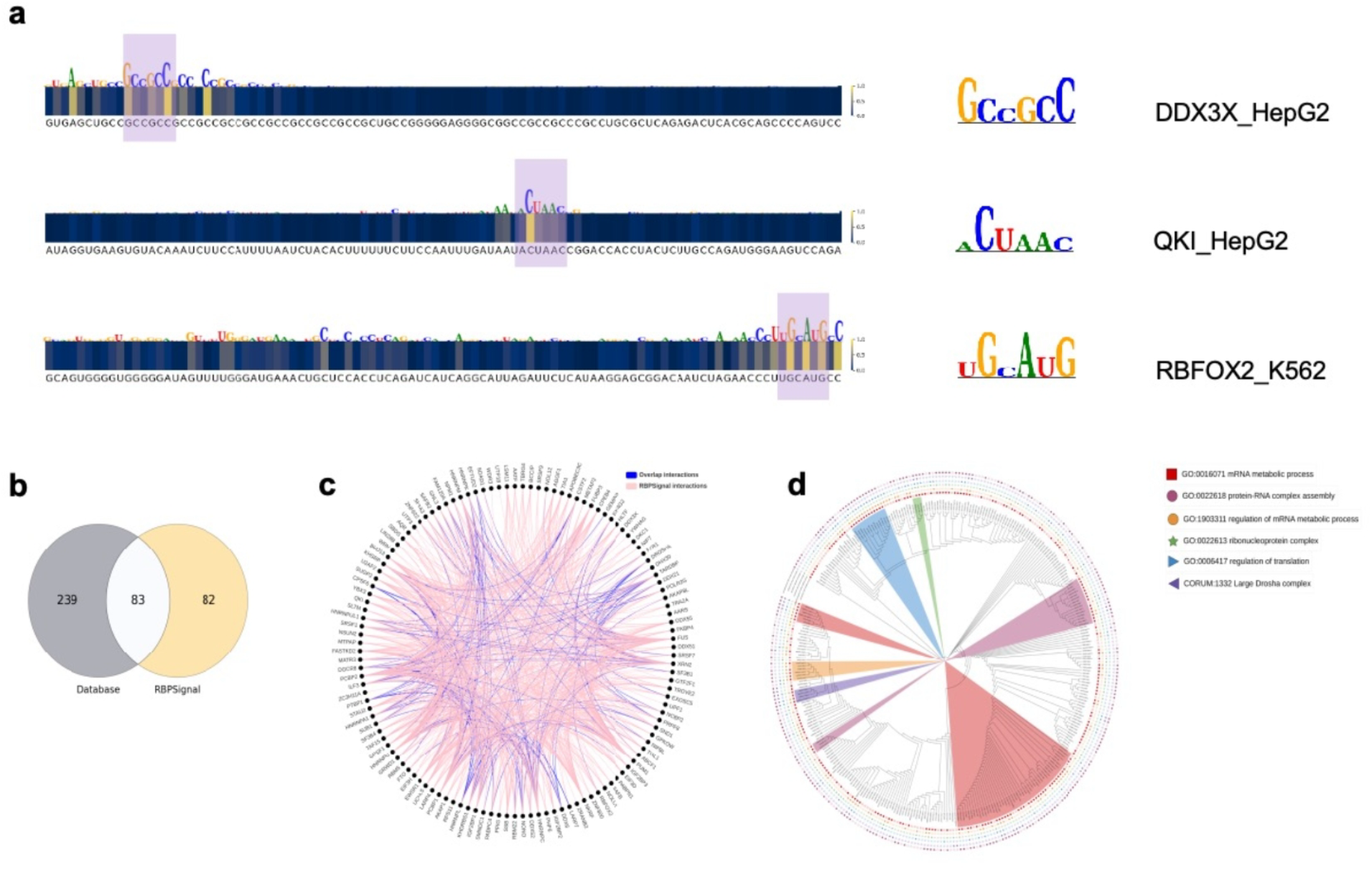
RBPSignal identifies effective motifs and offers biological insights. a. Motif identification using the Integrated Gradients interpretability method, demonstrated with examples of DD3X_HepG2, QKI_HepG2, and RBFOX2_K562. b. Venn diagram showing overlap between motifs identified by RBPSignal and known motifs in the database. c. Discovery of protein interactions using RBPSignal, with identified interactions compared against the STRING database. d. Phylogenetic analysis of proteins visualized as a circular tree, as discovered by RBPSignal.

To identify all potential motifs for each RBP We utilized IG across all RNA sequences in our dataset. For each sequence, the highest-scoring 7-mer bases were selected as the representative motif. Based on prediction values, we chose the top 40,000 sequences (or all sequences if fewer than 40,000 were available) and compared the motifs identified by RBPSignal to the motif reference database containing 322 known motifs, with some RBPs associated with multiple distinct motifs. Remarkably, 83 motifs overlapped between RBPSignal predictions and the database, as illustrated in Fig. 3b.

Given that multiple RBPs may share binding motifs, we hypothesized that these shared motifs could indicate potential interactions among RBPs. By filtering interactions based on q-values, we identified potential interactive relationships, represented by pink lines in Fig. 3c. Additionally, known interactions from the STRING database are depicted as blue lines in the same figure. Several predicted interactions aligned with previously reported interactions, which supports the biological plausibility of our motif-based interaction strategy. For instance, FUS and TAF15 are connected in our analysis, and previous studies have shown that FUS interacts with TAF15, and both proteins undergo phase separation in neurons(Kim, 2025). Similarly, HNRNPK and PCBP2 both involved in mRNA stabilization and translation regulation, protecting RNA from degradation(Smirnova et al., 2019).

Beyond protein-protein interactions, the motif analysis also suggests possible evolutionary connections among RBPs (Fig. 3d). RBPs sharing conserved motifs may belong to functional families and exhibit evolutionary relationships, highlighting their shared roles in RNA processing and regulation. These findings underscore the utility of RBPSignal in elucidating both functional and evolutionary insights into RBP-RNA interactions.

### Demonstration of RBPSignal for Viral RNA Analysis and Web Server Application

We validated the applicability of RBPSignal on the 5’UTR of the Zika virus (ZIKV) genome, with the sequence file downloaded from http://rasp.zhanglab.net/download/. Previous study has reported that the double-stranded RNA-binding protein SND1 interacts with ZIKV and influences its fitness(Zhang et al., 2022). Using RBPSignal, the original binding value was predicted to be 1.2639. However, after mutating a single nucleotide (G to U) in a highly contributive region, the binding value decreased to 0.9475, indicating a notable reduction in binding affinity. Another single-nucleotide mutation (U to C) in the same originally high-contributive region resulted in a predicted binding value of 1.0927. Although this was higher than that of the previous mutation, it still represented a decrease compared to the wild-type sequence, demonstrating RBPSignal’s sensitivity to sequence changes.

To facilitate broader research applications, we have developed a user-friendly web server that enables researchers to predict RBP-RNA binding signals. As illustrated in Fig. 4b, the web server provides visualization of binding signals, where peaks indicate potential binding regions, offering an intuitive tool for exploring RBP interactions. We analyzed the same transcript for three RBPs: AGGF1, FUBP3, and LARP7. The results demonstrate distinct variations in the distribution of binding signals among these proteins.

**Figure 4.**
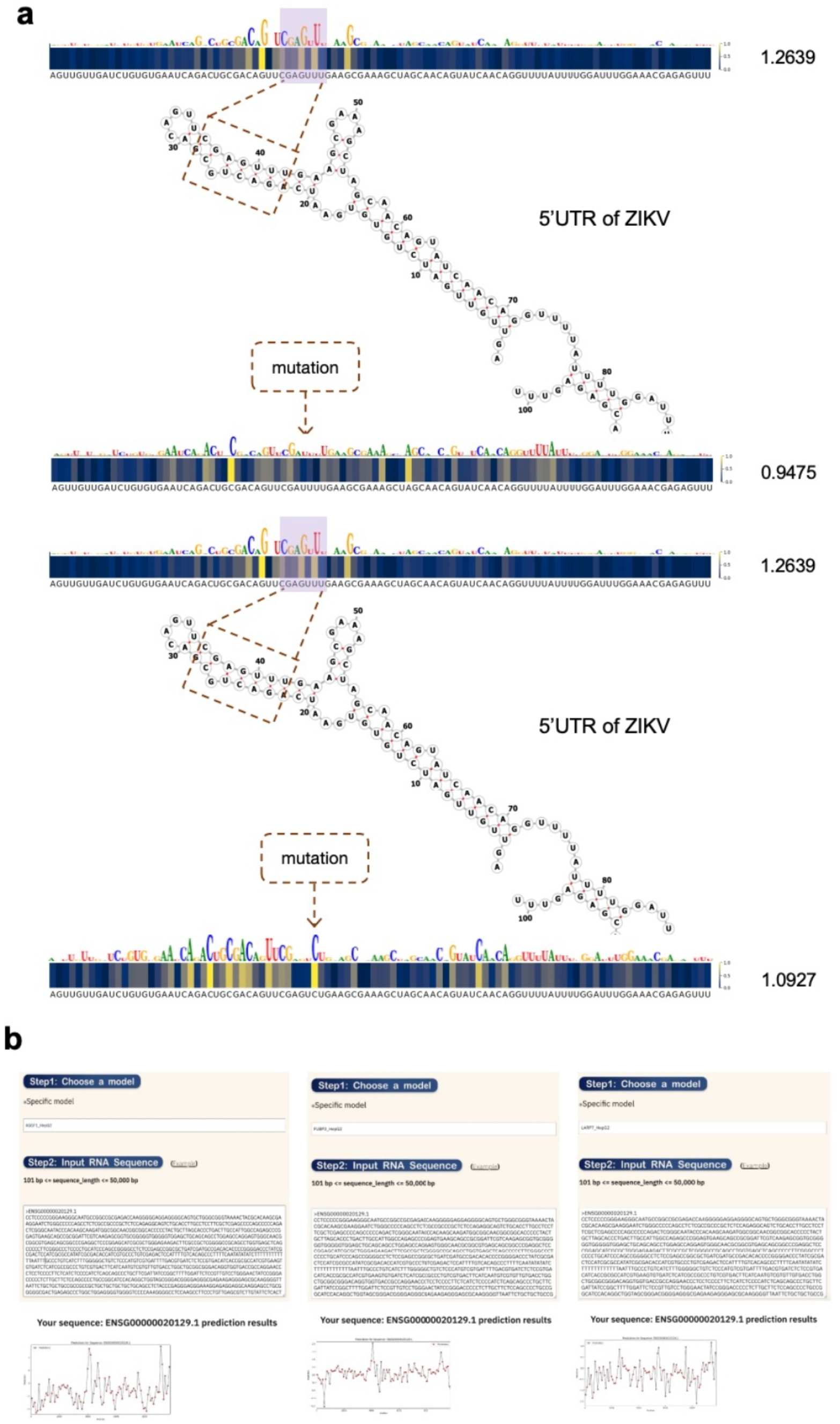
RBPSignal predicts specific RBP-virus interactions and provides a web server demonstration. a. Prediction results showing RBP SND1 binding to the ZIKV 5′ UTR with two different mutations. b. Example output from the RBPSignal web server, illustrating predicted binding signals.

## Discussion

RBPSignal represents an advancement in deciphering RNA-binding protein binding signals and brings biological interpretation into the underlying characteristics that regulate RBPs interactions with RNA. By leveraging a regression-based approach and deep learning networks, RBPSignal effectively bridges the gap between classification and regression tasks, capturing the complex signals associated with RBP binding and enabling a more nuanced understanding of how these proteins recognize and interact with their RNA targets.

From a biological perspective, RBPSignal illuminates the intricate mechanisms that RBPs interact with RNA, uncovering potential motifs that contribute to their functions in various cellular processes. Furthermore, RBPSignal explores protein evolution and interaction dynamics, providing valuable insights into the roles of RBPs in post-transcriptional regulation and their involvement in diverse biological pathways.

While RBPSignal has demonstrated promising results, there remains ample opportunities for improvement. Utilizing more advanced language models for sequence feature learning could further enhance its capabilities. Expanding the training and testing datasets to include a wider range of RNA sequences and RBP types would improve the model’s predictive accuracy and generalizability. Additionally, integrating multiple biological interpretability methods could provide deeper insights into the interaction mechanisms between RBPs and RNA. By improving these areas, future iterations of RBPSignal have the potential to advance our understanding of RNA-protein interactions and their impacts in health and disease.

## Data availability

165 out of 252 RBPs eCLIP data alignment files covering 124 specific RBPs in bam format are sourced from ENCODE eCLIP datasets. All processed data files are now available at https://github.com/lemonxh123/RBPSignal.

## Code availability

RBPSignal is now available at https://github.com/lemonxh123/RBPSignal with all packages implemented in Python. We have also provided a user-friendly server at http://www.csbio.sjtu.edu.cn/bioinf/RBPSignal/ for predicting RBP binding signals with RNA. Users can choose the precise RBP types and enter the query RNA sequences in the input box or upload a text file containing RNA sequences in FASTA format. Users can also download results directly from the web server.

## Author Contributions

Xiaoyong Pan and Xiaojian Liu conceptualized the project. Zidong Shu was responsible for dataset collection and preprocessing. The model was developed and trained by Xiaohan Ding, with methodological guidance provided by Shengfan Wang and Xiaojian Liu. Hong-Bin Shen and Xiaoyong Pan supervised and provided strategic direction for the project. Xiaohan Ding and Zidong Shu plot the figures and formatted them. Xiaohan Ding developed the online website and implemented its functionality. Xiaoyong Pan, Xiaohan Ding, Zidong Shu, Luhan Shi and Xiaojian Liu drafted the manuscript, with contributions from Yan Zhou, Biao Zhang and received constructive feedback from Hong-Bin Shen. All authors reviewed and approved the final manuscript.

### Funding

This work is supported by the National Natural Science Foundation of China (No. 62473257, 62201506), and the Science and Technology Commission of Shanghai Municipality (24ZR1435300, 24510714300).

### Competing interest

The authors declare no competing interests.

## Notes

### Competing Interest Statement

The authors have declared no competing interest.

